# NORMA-The network makeup artist: a web tool for network annotation visualization

**DOI:** 10.1101/2020.03.05.978585

**Authors:** Mikaela Koutrouli, Evangelos Karatzas, Katerina Papanikolopoulou, Georgios A. Pavlopoulos

## Abstract

NORMA is a web tool for interactive network annotation visualization and topological analysis, able to handle multiple networks and annotations simultaneously. Precalculated annotations (e.g. Gene Ontology/Pathway enrichment or clustering results) can be uploaded and visualized in a network either as colored pie-chart nodes or as color-filled convex hulls in a Venn-diagram-like style. In the case where no annotation exists, algorithms for automated community detection are offered. Users can adjust the network views using standard layout algorithms or allow NORMA to slightly modify them for visually better group separation. Once a network view is set, users can interactively select and highlight any group of interest in order to generate publication-ready figures. Briefly, with NORMA, users can encode three types of information simultaneously. These are: *i)* the network, *ii)* the communities or annotations and *iii)* node categories or expression values. Finally, NORMA offers basic topological analysis and direct topological comparison across any of the selected networks. NORMA service is available at: http://bib.fleming.gr:3838/NORMA or http://genomics-lab.fleming.gr:3838/NORMA. Code is available at: https://github.com/PavlopoulosLab/NORMA

## Introduction

In the biomedical field, networks are widely used to capture the interactions and the associations between bioentities (e.g. proteins, genes, small molecules, metabolites, ligands, diseases, drugs) in order to unravel how complex systems operate. Despite the great variety of visualization tools to efficiently represent such networks [1–6], visualization still remains a bottleneck as views often become incomprehensive due to the large number of nodes and edges. Therefore, ways to visually focus on the important parts of a network emerge. Popular state-of-the-art tools include the Cytoscape [7], Cytoscape.js [8], Gephi [9], Pajek [10], Ondex [11], Proviz [12], VisANT [13], Medusa [14], Osprey [15], Arena3D [16], Graphia (Kajeka) and BioLayout Express [17]. However, most of these tools focus on interactivity, layout and network exploration and lack efficiency in network annotation visualization.

ClueGO [18] for example, a Cytoscape plug-in [19], partially solves this problem by using pie-chart nodes to show groupings of nodes which share common Gene Ontology terms [20] or KEGG pathways [21]. Enrichment Map [22] on the other hand uses similar approaches to visualize gene-set enrichment results. STRING [23] uses pie-chart nodes to overlay Gene Ontology, KEGG [21] and Reactome [24] pathways on a network [25]. Osprey builds data-rich graphical representations from Gene Ontology (GO) annotated interaction data and is bound to the BioGRID repository [26]. NAViGaTOR [27], offers among others, a Gene Ontology annotated matrix view as well as a matrix layout to represent groupings. DyCoNet (28), a Gephi plugin, allows community detection and visualization. CellNetVis (29) is a web tool dedicated to the visualization of biological networks and the cellular components their nodes belong to. ClusterProfiler (31) is an R package that automates the process of biological-term classification and the enrichment analysis of gene clusters.

While many of these tools try to partially address the annotation visualization problem by mostly using node coloring or pie-chart nodes to show categories, they mainly focus on integrating networks with knowledge from existing repositories, require a steep learning curve to become familiar with, are bound to certain databases, or are offered in a command line version.

In this article, we present NORMA, a highly interactive web tool which is especially designed to visualize communities and network annotations using alternative representations. NORMA is designed for non-experts and is a tool of general purpose which makes it platform and concept independent; thus aiming to engage users from different fields.

## Materials and methods

NORMA is a handy web tool for interactive network annotation, visualization and topological analysis, and is able to handle multiple networks and annotations simultaneously. In its current version, the main interface is split into four Tabs namely: *i)* Upload, *ii)* Network, *ii)* Annotation and *iv)* Topology.

### The Upload Tab

Users can upload three different file types as input. These are *i)* the network file, *ii)* the annotation file and *iii)* the node coloring file.

The *network file* is an obligatory, 2-column (unweighted), tab-delimited file, containing all network connections of an undirected network. Notably, self-loops and multiple-edges are eliminated automatically. In its online version NORMA cannot accept a network of more than 10,000 edges.

The *annotation file* is an obligatory, 2-column, tab-delimited file which contains information about the defined groups. The first column contains the group names whereas the second column contains the node names in that group. This file may contain results from a functional enrichment (e.g. Gene Ontology or KEGG/Reactome pathways) or a clustering analysis (e.g. MCL output [28,29]: Group-x contains nodes a,b,c or Group-y contains nodes k,m,n).

The *node coloring file* is an optional, 2-column, tab-delimited file in which users may directly assign a node color to encode different types of information like for example gene expression values (e.g. red for a down-regulated gene or green for an up-regulated gene) or a node category (e.g. yellow for category-1 and blue for category-2). In this file, the first column contains the node names whereas the second column the nodes’ colors (e.g. red, green, yellow, blue, orange). Nodes without color assignment will be colored gray.

In NORMA, users can upload as many network and annotation files as they like. Every time a network or an annotation file is uploaded, a name can be given first. Once the network and annotation files have been named and uploaded, users have the option to visualize any network and overlay any of the selected annotations. Users can remove indifferent annotations or networks at any time. Network and annotation file contents are always shown as interactive tables in which one can search by suffix from the Search field.

### The Network Tab

This Tab consists of two sub-tabs dedicated to network analysis and visualization. These are: *i)* the Interactive Network and *ii)* the Automated Community Detection.

The *Interactive Network* sub-Tab offers a dynamic network visualization in its simplest form with the use of vizNetwork library. Nodes are connected with undirected edges whereas their coordinates are calculated using vizNetwork’s default force-directed layout. The network is fully interactive as zooming, dragging and panning are allowed either by using the mouse or the navigation buttons. Nodes can be selected and dragged anywhere on the plane, whereas the first neighbors of any node can be highlighted upon selection. The network view is automatically updated when a different network from the ones that have been uploaded is selected.

The *Automated Community Detection* Tab offers methods for automatic community detection. This is particularly important in cases where one does not have any external pre-calculated results to provide. The methods that are currently offered are:

- *Fast-Greedy* [30]: This function tries to find densely connected subgraphs (also called communities) via directly optimizing a modularity score.
- *Louvain* [31]: This function implements a multi-level modularity optimization algorithm for finding community structures and is based on the modularity measure and a hierarchical approach.
- *Label-Propagation* [32]: This is a fast, nearly linear time algorithm for detecting community structures in a network by labeling the vertices with unique labels and then updating them by majority voting in the neighborhood of the vertex.
- *Walktrap* [33]: This function tries to find densely connected subgraphs in a graph via random walks. The idea is that short random walks tend to restrict themselves in the same community.
- *Betweenness* [34]: Many networks consist of modules which are densely connected between themselves but sparsely connected to other modules. Clustering is made by ‘breaking’ the bridges which connect densely connected regions.

Once a community detection method has been selected, users can see the results as interactive and searchable tables or as static plots for an at-a-glance view. In order for users to take advantage of NORMA’s advanced interactive visualization capabilities, the automatically generated annotations must be first exported and then imported as annotation input files.

### The Annotation Tab

This Tab is NORMA’s strongest feature and is used to visualize annotated networks in an easy and user-friendly way. Annotated, are the networks which are accompanied by (pre-)defined clusters, communities, subgraphs, marked regions or neighborhoods. Here, users can select between any of the uploaded networks or annotation files and visualize them in combination (dropdown selection lists). This Tab consists of three sub-tabs. These are the: *i)* Convex Hulls, *ii)* Pie-chart nodes and *iii)* Venn diagrams.

#### Convex Hulls

In this Tab, the selected network is initially visualized after applying any of the offered layout algorithms and color-filled convex hulls are then used to highlight communities in a Venn-diagram-like view. A node might belong to more than one group. Groups are highlighted using visually distinct colors, whereas transparency is used to efficiently highlight the overlapping regions.

#### Pie-chart nodes

Here, the selected network is initially visualized after applying any of the offered layout algorithms and nodes are then visualized as pie-charts, divided into slices to illustrate the groups a node belongs to. If a node for example belongs to four groups, then the pie chart will consist of four equal slices colored with distinct colors. Nodes which do not belong to any group are marked gray.

#### Node coloring

Often, one might want to assign certain colors to nodes in order to encode certain types of information. In a gene expression network for example, one might want to highlight the up- and down-regulated genes. Once an expression file has been uploaded, nodes in the Convex Hull Tab will be filled with the color of interest whereas nodes in the Pie-Chart Tab will appear with a colored border. As node coloring is an optional feature, one can enable or disable this functionality at any time (selection box).

To briefly demonstrate this functionality, we queried the STRING database for the *BCAR3* Human gene and retrieved its interactions. We used all STRING channels (Text-mining, Experiments, Databases. Co‑expression, Neighborhood, Gene Fusion, Co‑occurrence) using the medium confidence threshold (0.4) and allowing no more than 10 interactions for the first interactors. After exporting STRING’s KEGG-analysis results and importing them into NORMA application, we highlighted four pathways. These are the protein processing in endoplasmic reticulum (genes involved: DERL1, DERL2, NPLOC4, NSFL1C, SYVN1, UFD1L, VCP, VIMP), the Regulation of actin cytoskeleton (genes involved: BAIAP2, BCAR1, CDC42, PAK1, PAK2, PXN, WAS, WASL), the Axon guidance (genes involved: CDC42, PAK1, PAK2, PARD6A, PARD6B) and the Bacterial invasion of epithelial cells (genes involved: BCAR1, CDC42, PXN, WAS, WASL). The network along with the four selected pathways is shown in Figure 1.

**Figure 1.**
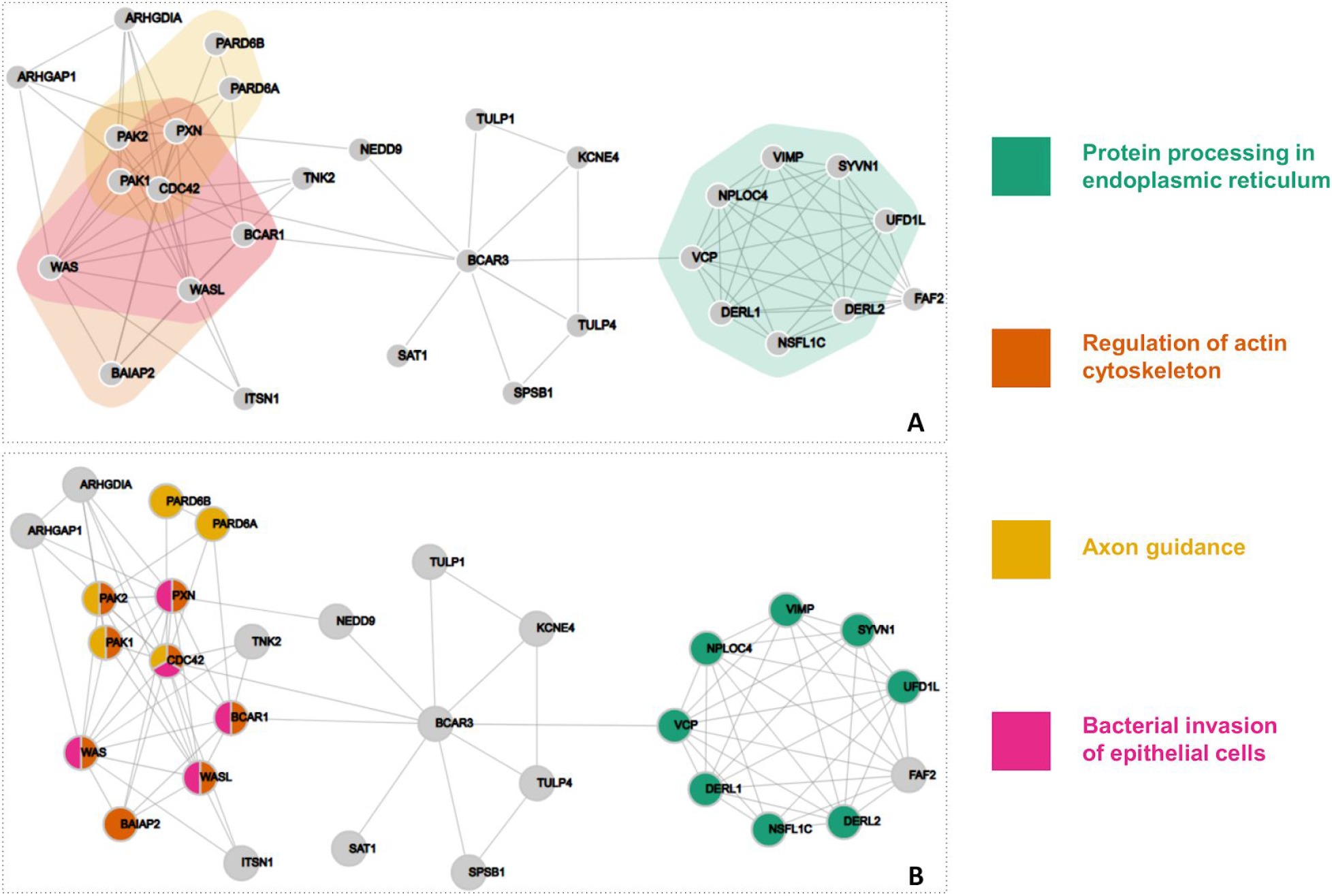
Visualization of BCAR3 interactors from STRING with four highlighted KEGG pathways. A) Color-filled convex-hulls. B) Pie-chart nodes.

##### Layouts

Users, can choose between several layout algorithms to visualize a network. In its current version, NORMA offers the layout algorithms offered by the igraph library [35]. These are the:

- *Fruchterman-Reingold* [36]: It places nodes on the plane using the force-directed layout algorithm developed by Fruchterman and Reingold.
- *Random*: This function places the vertices of the graph on a 2D plane uniformly using random coordinates.
- *Circle*: It places vertices on a circle, ordered by their vertex ids.
- *Kamada-Kawai* [37]: This layout places the vertices on a 2D plane by simulating a physical model of springs.
- *Reingold-Tilford* [38]: This is a tree-like layout and is suitable for trees or graphs without many cycles.
- *LGL*: A force directed layout suitable for larger graphs.
- *Graphopt*: A force-directed layout algorithm, which scales relatively well to large graphs.
- *Gem* [39]: It places vertices on the plane using the GEM force-directed layout algorithm.
- *Star*: It places vertices of a graph on the plane, according to the simulated annealing algorithm by Davidson and Harel.
- *Grid*: This layout places vertices on a rectangular 2D grid.

For better clarity, NORMA gives the option to users to slightly modify the selected layout in order to make groups as distinct as possible, thus avoiding unnecessary overlaps which may occur due to the original layout. To do this, NORMA starts by introducing virtual nodes (one per group) and connects them with the nodes this group relates to by assigning a heavy weight. This way, the virtual nodes will pull together the nodes they relate to regardless of the selected layout. Upon completion, virtual nodes are then removed. The whole process is schematically shown in Figure 2.

**Figure 2.**
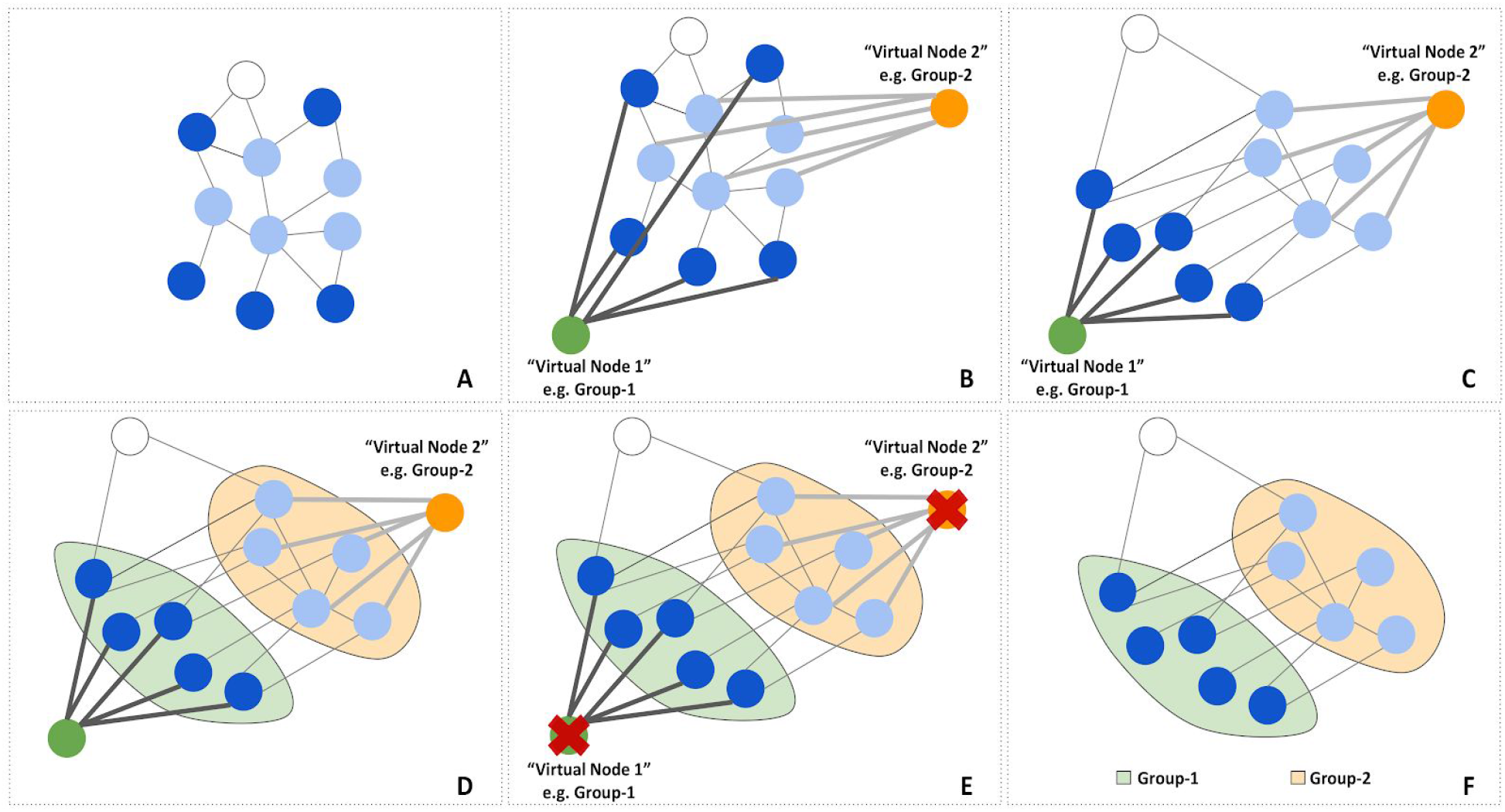
NORMA’s layout process. A) Norma starts with a network using random coordinates. B) Virtual nodes are introduced (one per group) and are connected with heavy weights to the nodes they relate to. C) A selected igraph layout is applied including the virtual nodes. Virtual nodes are expected to pull together the nodes they relate to due to the heavy weight. D) Once the selected layout algorithm is finished, the color-filled convex hulls are introduced. E) Virtual nodes are removed. F) The final network view.

#### Interactivity and Visualization

NORMA gives a variety of options for the creation of optimal custom views. Network zoom in/out and panning functionalities are offered while users can interactively drag any node and place it anywhere on the plane. In addition to the visualized networks, groups are shown in an interactive table whose rows are colored accordingly. By selecting one or more groups, one can show the convex hulls as well as the pie-chart nodes of interest. Colored groups (rows) in the table correspond to colored groups in the offered views and vice versa. In addition, users have the option to show and hide the labels or only keep the labels of the selected groups of interest while labels below a certain zoom level are hidden for clarity. Finally, sliders to adjust node and label sizes as well as a slider to scale the network size are offered.

#### Venn Diagrams

Users are allowed to choose between any pair of nodes and visualize the common groups they belong both as a Venn diagram and as a table. As it is not in the scope of NORMA to provide more complex Venn diagrams, users are encouraged to visit other online applications dedicated to this purpose [40–43].

### The Topology Tab

Inspired by NAP’s [44] and Cytoscape’s topological network analysis features, in this Tab, NORMA offers an automated topological analysis and direct comparison of topological features between two or more selected networks. The topological measured that are offered by the igraph library are the:

- *Number of Edges*: Shows the number of connections in the network.
- *Number of Nodes*: Shows the number of nodes in the network.
- *Density*: The density of a graph is the ratio of the number of edges and the number of possible edges.
- *Average path lengt*h: The average number of steps needed to go from one node to another.
- *Clustering Coefficient*: A metric which shows if the network has the tendency to form clusters. Values between 0 and 1.
- *Modularity*: This function calculates how modular is a given division of a graph into subgraphs.
- *Average Eccentricity*: The distance from a particular vertex to all other vertices in the graph is taken and among those distances, the eccentricity is the highest of distances.
- *Average number of Neighbors*: It is the total number of neighbors per node divided by the number of nodes.
- *Centralization betweenness*: It is an indicator of the average centrality in a network. It is equal to the number of shortest paths from all vertices to all others that pass through that node. Betweenness centrality quantifies the number of times a node acts as a bridge along the shortest path between two other nodes.
- *Centralization degree*: It is defined as the average value of the number of links incident upon a node, across all nodes of a network.

The Topology Tab is divided into two sub-tabs. These are: *i)* the Summaries and *ii)* the Comparative Plots. The *Summaries* Tab shows the aforementioned topological measures in a numerical form as table. Users can select one or more topological measures of interest and expand the table accordingly. Notably, this can be done for one network at a time upon selection (dropdown selection list). In the *Comparative Plots* Tab, users can directly compare the topological features of two or more networks simultaneously. In contrast to the previous Tab, users are allowed to select one topological feature at a time (radio buttons) but as many networks as they like (check boxes). Once two or more networks and one topological feature have been selected, direct comparisons can be made by the generated bar charts. A slider to adjust the chart height is offered.

### Implementation

NORMA is implemented in R and its interface in Shiny. Network analysis features are mainly based on the igraph library. Network interactivity and visualization are implemented in d3.js. NORMA service is available at: http://bib.fleming.gr:3838/NORMA or http://genomics-lab.fleming.gr:3838/NORMA Code is available at: https://github.com/PavlopoulosLab/NORMA

### A general overview

In order to summarize NORMA’s functionality, we briefly show its features in Figure 3. The network shown here has been generated by querying STRING for human TP53 protein (default settings). Figure 3A shows the unweighted network in its simplest form. Figure 3B shows a static version of the same network clustered with NORMA’s Louvain clustering algorithm. Figure 3C shows the network and its groups (manually and randomly assigned) both as color-filled convex hulls and as pie-chart nodes. Groups are also shown as a table (legend) and colors between the network views and the table are consistent. In both views, node coloring encodes additional information (full color in the case of convex-hulls or border color in the case of pie-chart nodes). Figure 3D shows a Venn diagram to highlight the common groups between two selected nodes and Figure 3E shows the topological network analysis of the TP53 network and a direct comparison (e.g. number of edges) with the BCARC3 network which was described previously in Figure 1.

**Figure 3.**
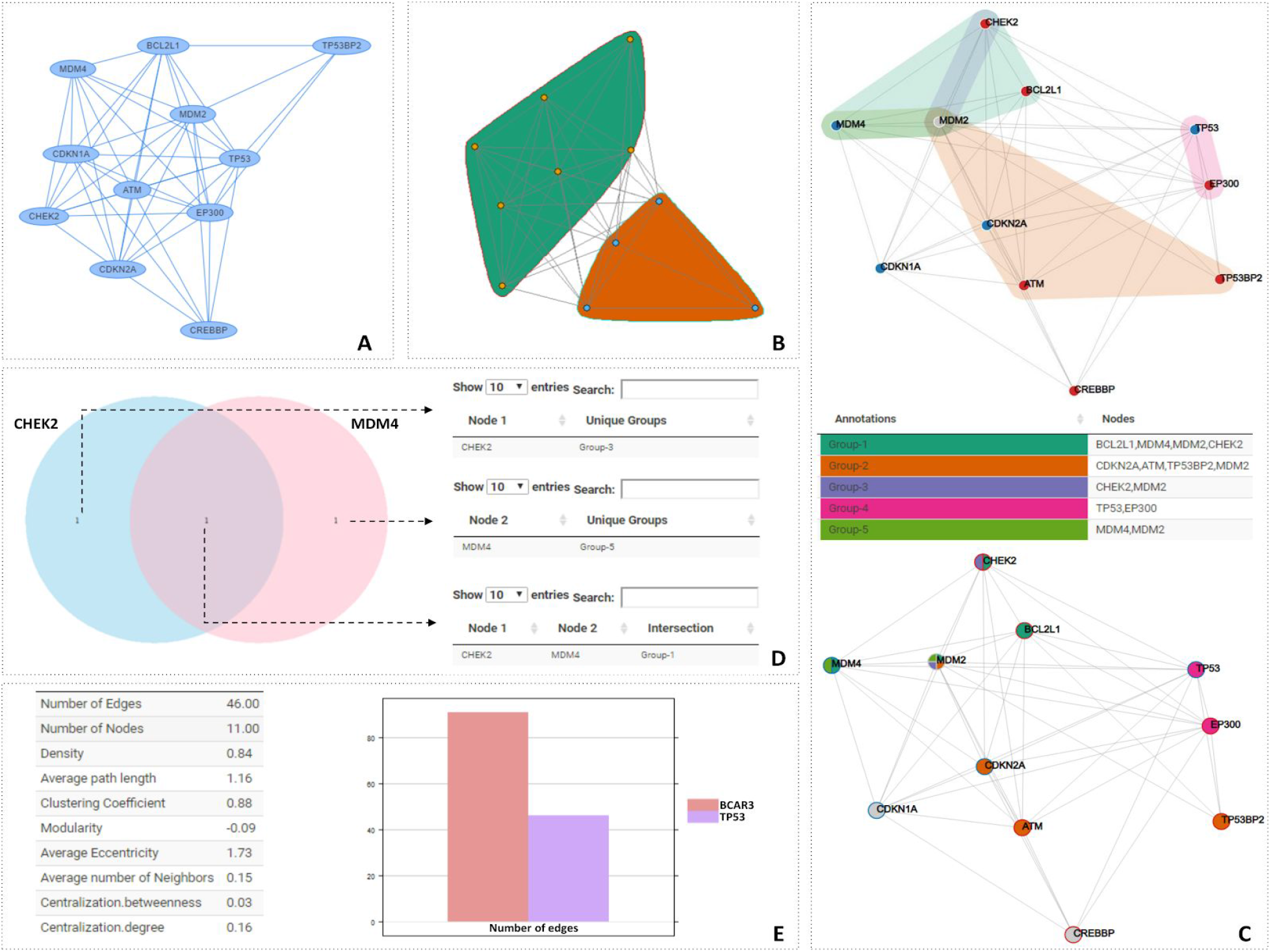
NORMA’s overview. A) The TP53 network. B) Community detection using the Louvain algorithm. C) *Upper part*: Convex hulls with colored nodes, *Lower part*: Pie-chart nodes with border colors. *Middle part*: Groups visualized in an interactive table (group selection/isolation is allowed). D) Venn diagrams to show common groups between any pair of selected nodes. E) *Left part*: Basic topological analysis of the TP53 network, *Right part*: bar chart to directly compare one topological feature between two networks.

## Results

In order to demonstrate NORMA’s capabilities in a real case study, we used the experimental data described in [45]. Null-Tau mutants in Drosophila were used to demonstrate broad changes in their brain proteome with mass-spectrometry. To briefly describe the protocol used in this study, tauKO mutant flies [46] and Cantonised-w1118 control flies were cultured on standard wheat-flour-sugar food supplemented with soy flour and CaCl2, at 25°C in 50–70% relative humidity in a 12 h light/dark cycle. Three to four biological and two technical replicas from each genotype were used with each biological replica counting 10 brains. The samples were digested following the filter-aided sample preparation (FASP) method using spin filter devices with 10kDa cutoff (Sartorius, VN01H02) [47]. The peptide products were detected by an LTQ Orbitrap XL mass spectrometer (Thermo Fisher Scientific, Waltham, MA, U.S.A.) as described elsewhere [48]. The mass spectral files (raw files) were processed using MaxQuant software (version 1.5.3.30) [49] against the complete Uniprot proteome of Drosophila melanogaster (Downloaded 1 April 2016/ 42456 entries) and a common contaminants database by the Andromeda search engine. For the calculation of the protein abundances, label-free quantification (LFQ) was performed with both “second peptides” and “match between run” options enabled. Statistical analysis was performed using Perseus (version 1.5.3.2).The label free quantified proteins were subjected to statistical analysis with student’s t-test (permutation-based p-value with 0.05 cutoff).

Starting from the identified 503 differentially expressed proteins, we used STRING to extract the connections among 358 of them and performed STRING’s KEGG analysis to highlight several important pathways (Figure 4A). Up- and down-regulated proteins are marked in green and red accordingly. The most affected pathways upon dTau loss in the central brain are shown as color-filled convex hulls and involve molecular processes reported altered in Tauopathies. Tau for example, is known to bind to ribosomes in the brain and impair their function reducing protein synthesis [50], an effect enhanced in Tauopathy brains [51]. In agreement, loss of dTau, resulted in elevated translation and protein synthesis as reflected by the increased expression of translation machinery components, including initiation factors, proteins which control gene expression or silencing and ribosomal proteins. Spliceosome disruption and altered pre-mRNA processing are also emerging as potential contributors to Tau-mediated neurodegeneration [52]. On the other hand, transgenic Tau elevation by expression of human Tau in the fly CNS, triggered the reduction of multiple spliceosomal proteins [53]. In accordance with these results, several RNA-binding proteins and spliceosome components are more abundant in the mutant. Congruently with translation, proteins mediating proteasomal catabolism are equally elevated. Interestingly, proteasome activation can be viewed as an adaptive response to support elevated protein synthesis by increasing the size of the intracellular amino acid pool [54]. Finally, proteins implicated in energy-metabolism (oxido-reductases) and responsiveness to oxidative stress, appear as differentially regulated in the mutants, with about half of them attenuated by its loss. This is consistent with the reported accumulation of oxidative damage markers in Tauopathy patients [55] and the effect of oxidative stress in mediating Tau-induced neurotoxicity in flies [56].

**Figure 4.**
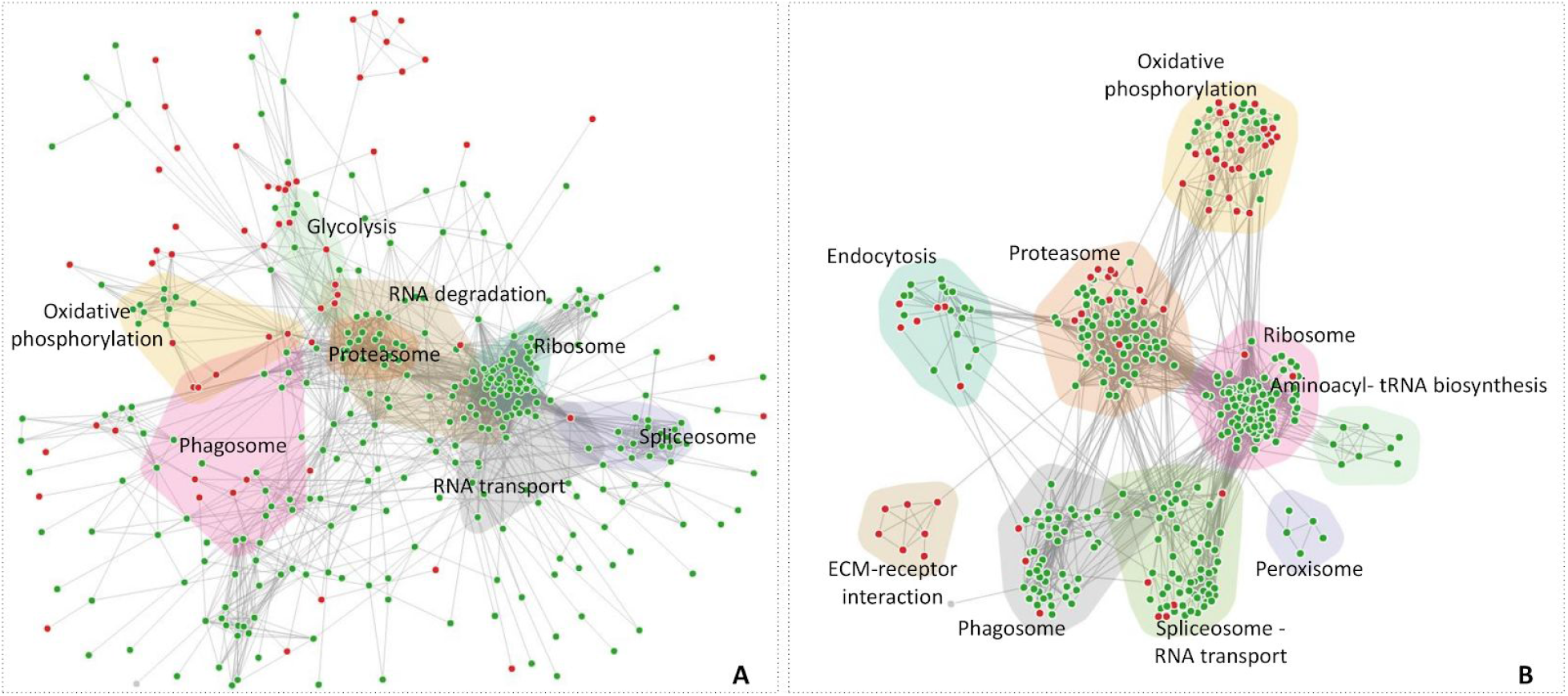
Drosophila’s Tau interaction network. A) 8 major KEGG pathways highlighted. B) Louvain clustering and functional annotation with DAVID application.

To partially regenerate the annotation in Figure 4A, we used NORMA’s Louvain community detection algorithm to cluster the network based on its topology only and DAVID [57] application to perform pathway and GO enrichment analysis. In Figure 4B, results revealed nine distinct communities which are highlighted. Within these communities, 4/22 proteins are involved in endocytosis, 21/75 in proteasome, 3/5 in peroxisome, 43/85 in ribosome, 15/58 in spliceosome, 15/49 in oxidative phosphorylation, 4/12 in ECM receptor, 8/47 in phagosome and 7/8 in aminoacyl -tRNA biosynthesis.

## Discussion

NORMA is a web tool for network annotation visualization. It is able to handle multiple annotations and networks simultaneously and is made to aid researchers in producing high-quality publication-ready figures as well as to explore novel hypotheses. In its current version, NORMA is not bound to a certain database or repository while in the future it will offer alternative visualization options (e.g. Annotated Matrix View, Hive-Plots, Circos-like diagrams with edge bundling support). We believe that due to its interactivity and ease-of-use, NORMA is a handy tool for selective annotation visualization designed for non-experts without prior familiarity with command line tools or scripting.

## Funding

GAP and MK were supported by the Operational Program Competitiveness, Entrepreneurship and Innovation, NSRF 2014-2020, Action code: MIS 5002562, co-financed by Greece and the European Union (European Regional Development Fund). GAP was also supported by the Hellenic Foundation for Research and Innovation (H.F.R.I) under the “First Call for H.F.R.I Research Projects to support Faculty members and Researchers and the procurement of high-cost research equipment grant”. EK has been supported by the Action Strengthening Human Resources, Education and Lifelong Learning, 2014–2020, co-funded by the European Social Fund (ESF) and the Greek State (MIS 5000432). KP was supported by a grant from the Stavros Niarchos Foundation to the Biomedical Sciences Research Center “Alexander Fleming,” as part of the initiative of the Foundation to support the Greek research center ecosystem.

## Acknowledgements

We thank Dr. Yorgos Sofianatos for his valuable feedback and Dr. Alexandros Dimopoulos for the server setup.

## Authors’ contributions

MK was the main developer of the tool. EK worked on network interactive visualization. KP provided data and wrote the result section. GAP supervised the whole project. All authors wrote parts of the manuscript and have approved its final version.

## Conflict of interest

Authors declare that there is no conflict of interest.

